# Improving real-time search through deep learning-enhanced fragment ion indexing

**DOI:** 10.64898/2026.10.27.762387

**Authors:** Jimmy K. Eng, Michael R. Hoopmann, Devin K. Schweppe, Michael J. MacCoss, William S. Noble, Bo Wen

**Affiliations:** Department of Genome Sciences, University of Washington; Proteomics Resource, University of Washington; Paul G. Allen School of Computer Science and Engineering, University of Washington

**Keywords:** fragment ion indexing, database search, real-time search, mass spectrometry, proteomics, deep learning

## Abstract

The fragment ion indexed database searching approach enables highly efficient tandem mass spectrometry database searches, including real-time search workflows. However, the index can require tens of gigabytes of memory, often exceeding the free memory available on instrument acquisition computers running the search. This restricts the size of the searchable database, the variable modification space, and the number of searches that can run concurrently. To address this problem, we developed an index-pruning pipeline within the Comet database search tool using Carafe, a deep learning tool that predicts fragment ion intensities based on peptide sequences. In our approach, only fragment ion positions that are predicted to be observable are written in the index, eliminating entries that are unlikely to contribute to spectrum detection. We evaluated this approach on two search spaces in three index configurations to demonstrate that masking significantly reduced the number of fragment ion index entries and, consequently, peak search memory use. Masked searches were also faster than unmasked searches and identified a comparable number of peptide-spectrum matches. Because all prediction and masking steps are performed before searching, their cost is amortized across repeated, future searches of the same database and modification configuration, introducing no per-spectrum search overhead. The complete implementation is open source with an Apache license.

## Introduction

Tandem mass spectrometry (MS/MS) is the leading technology for characterizing proteomes, with applications such as the analysis of post-translational modification signaling, mapping protein–protein interaction networks, biomarker discovery, and drug target identification. ^1^ In data-dependent acquisition (DDA), the instrument acquires a precursor survey scan (MS1), selects the most abundant precursors, and isolates and fragments those precursors to produce fragmentation scans (MS2). Decisions about which precursors to fragment, or which dissociation technique to use, must typically be made without knowing which peptides the precursors represent, frequently leading to suboptimal choices by the instrument control software.

Real-time search (RTS) addresses this limitation by identifying each MS2 spectrum within the duty cycle in which it is acquired, so that subsequent instrument decisions can be based on the identified precursor. ^2–4^ In isobaric labeling-based protein quantification, restricting the accurate but time-consuming synchronous precursor selection MS3 scan (SPS-MS3) to spectra that yield a confident peptide-spectrum match (PSM) allows RTS-MS3 to quantify a comparable number of proteins in roughly half the instrument time required by conventional SPS-MS3. ^2–4^ The same principle improves proteome coverage in demanding applications such as single-cell multiplexed proteomics. ^5^ RTS imposes strict time requirements: spectrum processing, candidate generation, scoring, and thresholding must all finish within the time window in which a peptide ion elutes from the column. Because tens to hundreds of peptide ions trigger MS/MS acquisitions concurrently, the practical per-spectrum time budget is in the millisecond range.

The Orbiter platform ^4^ addresses this requirement by incorporating the open source search engine Comet ^6,7^ directly into the instrument control loop. To increase search speed further, Comet was extended with a fragment ion index (Comet-FI) search mode, ^8^ adopting the inverted index approach introduced to proteomics by MSFragger ^9^ and since implemented in MetaMorpheus, ^10^ Sage, ^11^ and Telescope ^12^. Comet-FI precomputes the theoretical b-ions and y-ions of every candidate peptidoform and stores them in an inverted index that maps each binned fragment mass to the peptidoforms that produce it. Querying the observed fragment peaks against this index yields the peptidoforms that share at least a minimum number of matched fragments, and only those candidates proceed to full scoring. This filtering step preserves detection accuracy while reducing the median single-spectrum RTS times.

The speed gain of fragment ion indexing comes at a memory cost, because the index must be held in memory for the duration of the search. Unlike offline searches, in which the index can be loaded and searched in subset batches, RTS has no practical way to load only part of the search space. Memory use grows further when variable modifications are enabled, because every modified form of a peptide contributes a complete ion series; broad peptide length ranges, relaxed enzymatic constraints or combinatorial modifications can produce indexes of many tens of gigabytes. This is particularly problematic because RTS typically runs on the instrument control computer. Thermo instrument control systems commonly ship with 32 GB of random access memory, much of it already occupied by the operating system and acquisition software, leaving well under 15 GB for running other applications such as searching. Furthermore, in order to perform RTS on acquisition methods that interleave dissociation types (CID, HCD, EThcD, etc.), a separate index for each dissociation method is required to be resident in memory simultaneously. Shrinking the fragment ion index is therefore essential for supporting large search spaces and concurrent RTS workflows on resource-constrained hardware.

In this work, we describe a strategy to reduce the memory footprint of the inverted index. The strategy makes use of the fact that peptide fragmentation is highly non-uniform: only a subset of the theoretically possible fragment ions, and an even smaller subset of their neutral loss (NL) variants, are observed at significant intensity. Furthermore, deep learning-based fragment intensity predictors are now accurate enough to be used routinely for re-scoring search results and generating *in silico* spectral libraries. ^13–17^ We therefore hypothesized that predicted intensities could be used to trim the fragment ion index by discarding fragments that are unlikely to be observed. Accordingly, we describe an extension of Comet-FI built on the fragment intensity models in Carafe. ^16,17^ For a given protein database and modification set, every candidate peptidoform is exported from Comet and analyzed by Carafe, and the predictions are converted into a compact binary mask of the fragment ion positions expected to be observable (Figure 1, bottom). Carafe can also predict modification-specific neutral loss species, such as the dominant phosphopeptide neutral loss, which Comet’s fragment ion index did not previously support. Because the search space is fixed before the experiment, all predictions can be computed in advance at no cost to acquisition-time RTS. We demonstrate that this approach substantially reduces the fragment ion index size and peak memory use, increases search throughput, and leaves the number of confident PSMs accepted at a 1% FDR threshold essentially unchanged, within *−*1.3% to +2.4% of the unmasked count.

**Figure 1:**
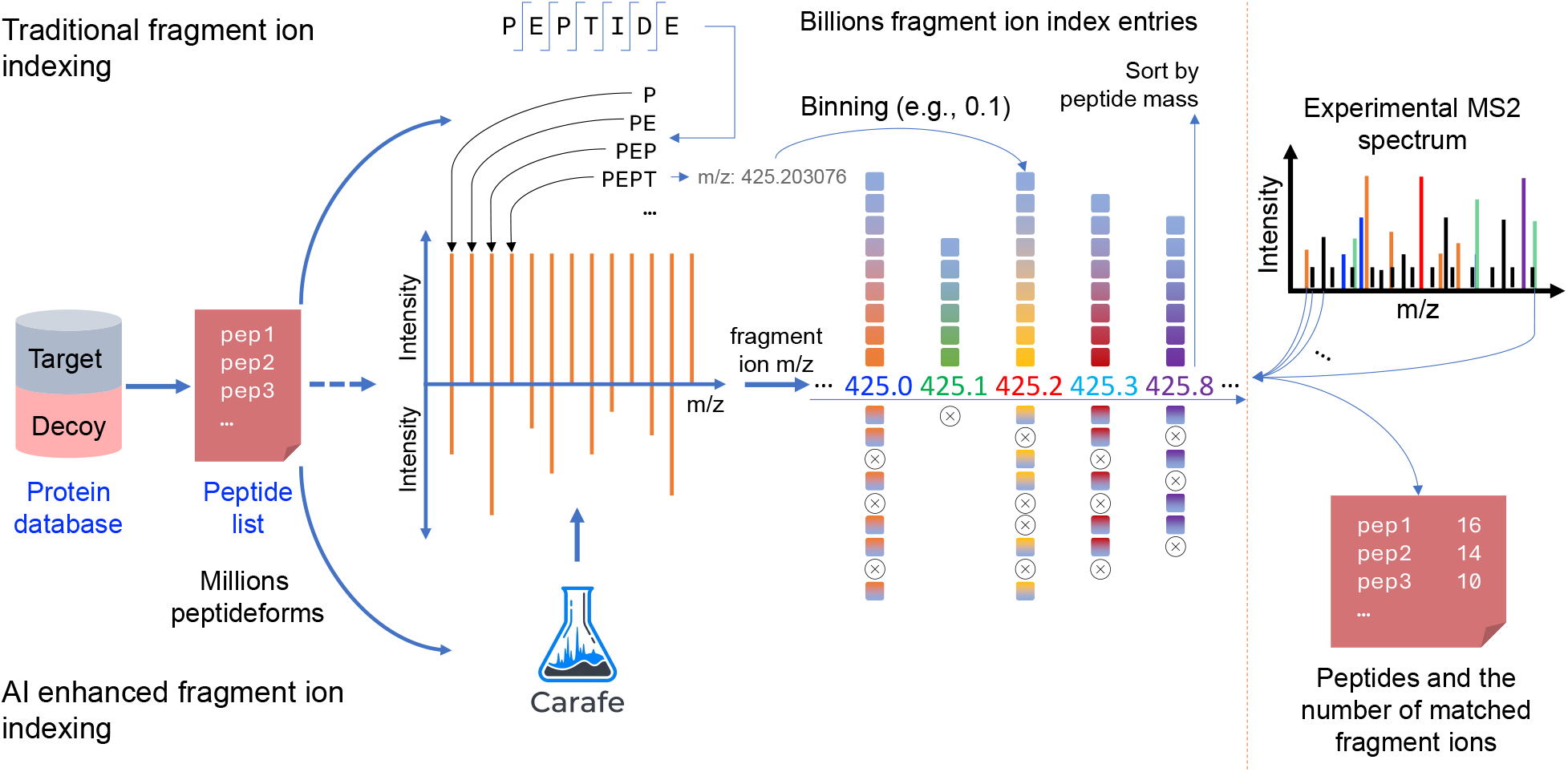
Overview of traditional versus AI-enhanced fragment ion indexing in Comet. Millions of candidate peptidoforms are generated from a target-decoy protein database and passed to Comet-FI. In traditional fragment ion indexing (top), each peptidoform (e.g., PEPTIDEK) is exhaustively fragmented into its complete b-ion/y-ion series (P, PE, PEP, PEPT, …), and every resulting fragment m/z is binned (e.g., 0.1 Da) into an inverted index mapping each binned fragment mass to the peptidoforms that produce it, yielding billions of fragment ion/peptide pointers sorted by peptide mass. In AI-enhanced fragment ion indexing (bottom), the same candidate peptidoforms are first passed through Carafe, a deep learning fragment ion intensity predictor, and only fragment ion positions predicted to be observable are written into the index. At search time (right), an experimental MS/MS spectrum is queried against the index to retrieve, for each candidate peptide, the matched fragment ions used for downstream scoring.

## 1 Materials and Methods

### 1.1 Software implementation

The feature was implemented in the Comet C++ code base (fragment ion index creation and search) and a set of standalone Python utilities that translate between Comet’s on-disk index format and Carafe’s input/output conventions. Comet itself does not invoke Carafe or perform machine learning inference during search execution; all inference is performed ahead of time. The Comet project’s C# RealtimeSearch example application for RTS integration was used to evaluate the masking performance.

### 1.2 Fragment ion mask generation

The fragment mask is generated once for a given protein database and variable modification configuration (Figure 1) by a series of resumable stages exposed through a single Python command line interface that runs identically on Windows and Linux. Individual stages are subcommands that execute the complete workflow. (1) Comet digests the database and builds a plain peptide index of unmodified peptides and their protein references. All variants that would later be enumerated during index construction, including every combination of variable and terminal modifications and each charge state, are then exported with metadata linking each variant to its index entry. (2) A standalone Python implementation of Comet’s plain peptide index reader then converts the exported variants to the AlphaBase sequence/modification/charge representation used by Carafe, and writes a mapping file that preserves the correspondence to Comet’s variants. (3) Carafe predicts fragment ion intensities across charge states using pretrained models without experiment-specific fine-tuning. In phosphorylation mode, Carafe uses the pretrained phosphorylation model from AlphaPeptDeep ^15^ and supports phosphorylation-specific neutral loss ions. For TMTpro peptides, Carafe’s bundled TMT CID model was used: a fine-tuning of its TMT HCD model on 5.7 million iontrap CID spectra of TMT6plex-labelled peptides (training pipeline and evaluation in the Carafe repository, `tools/tmt/ms2cid`, manuscript under preparation). The input is partitioned into fixed-size chunks processed by independent Carafe instances through a wrapper script; progress is saved after every chunk so that an interrupted run resumes without the need to recompute completed predictions. (4) The predictions are then converted to a compact binary prediction store (.cps). The raw Carafe parquet output for the phosphoproteomic search space evaluated here (215.7 million peptidoform-charge rows) is 78 GB and includes data currently not used for masking, such as predicted retention times. The .cps format retains only the fragment ion intensities, neutral loss shifted intensities, and a base peak reference intensity per variant, quantized to two bytes per value, and is 50.3 GB for that search space. (5) A final tool generates the mask. For each variant, the fragment ladder positions of the base and neutral loss series are evaluated independently, and a position is retained if its predicted intensity is at least a user-defined fraction of the variant’s predicted base peak intensity; all masks in this work used 10%. A minimum-kept-peaks parameter retains the most intense positions whenever fewer than the specified number would otherwise be kept, preventing excessive pruning; all masks in this work kept at least six positions per variant. These two settings were held fixed, and their effect on index size and detections was not explored here.

The mask is written as a binary .fi_mask file with a header (source index fingerprint, peptide count, and modification configuration) followed by a fixed-length record per variant holding four 64-bit bitmasks for retained b-ion, y-ion, neutral loss b-ion, and neutral loss y-ion positions, sorted for binary lookup. A single prediction store serves every index configuration that enumerates the same peptidoform variants, which the pipeline verifies before building a mask: for example, masks for neutral loss enabled and disabled indexes were both derived from one Carafe pass.

### 1.3 Search time masking

At search time, either during a batch Comet-FI search or an RTS session, the mask file is loaded once after a plain peptide index is read and before the in-memory fragment ion index is built. The mask’s embedded content fingerprint, peptide count, and variable modification configuration are validated against the active peptide index and search parameters, and any mismatch terminates the search so that a mask built for a different index or modification configuration can never be applied. Once a mask is accepted, each candidate b-ion, y-ion and, where applicable, neutral loss ion is evaluated against it during index construction, and only positions predicted to be observable are inserted. A variant that has no entry in the accepted mask, for example because no prediction was produced for it, is indexed without filtering, so an incomplete mask can reduce filtering efficiency but never removes candidates from a search. Masking does not change the set of peptidoforms in the index; it prunes only the entries of retained variants, acting as a preprocessing filter on the candidate fragment pool.

### 1.4 Datasets and search configurations

Two datasets, each defining one search space, were evaluated in three index configurations: the TMTpro search space, and the phosphoproteomic search space indexed with and without neutral loss ions. The first dataset was generated from a TMTpro 16-plex experiment and was downloaded from PRIDE with accession number PXD043660. Specifically, these data come from a TMTpro 16-plex sample composed of four human cell lines, acquired with a real-time search MS3 method on a Thermo Orbitrap Ascend mass spectrometer, in which MS2 spectra are acquired by CID in the ion trap (NCE 34) and reporter ions are measured in subsequent SPS-MS3 scans ^18^. Two basic-pH fractions (A2 and A4; files `Ascend_A2_65min_4cell.raw` and `Ascend_A4_65min_4cell.raw`) were searched. The TMTpro search used a tryptic digest with up to two missed cleavages, peptide length 7–50, static modifications of carbamidomethyl cysteine and TMTpro (+304.207146 Da) on lysine and every peptide N-terminus, variable oxidized methionine, a 20 ppm precursor tolerance with 0/1/2 ^13^C isotope offsets, and ion trap fragment settings. The second dataset was generated from a phosphoproteomic experiment and was downloaded from PRIDE with accession PXD011175. These data are derived from a phosphorylation sample composed of IMAC-enriched human malignant melanoma lymph node metastasis tissue analyzed on a Thermo Q Exactive HF-X Orbitrap mass spectrometer (HCD, NCE 25, MS2 in the Orbitrap at 60,000 resolution) ^19^; two biological replicate acquisitions (R1 and R2; files `MM2_R1.raw` and `MM2_R2.raw`) were searched. These searches used a tryptic digest with up to two missed cleavages, peptide length 7–40, static carbamidomethyl cysteine, variable oxidized methionine and phospho serine, threonine, and tyrosine, a 20 ppm precursor tolerance with 0/1/2 ^13^C isotope offsets, and highresolution fragment settings. This sample was searched both without and with the 97.9769 Da phosphate neutral loss fragments indexed (“no NL” and “with NL” in Figures 2 and 3). All searches used the UniProt ^20^ human canonical proteome, with common contaminants, and reversed decoys (20,454 target and 20,454 decoy sequences). Detailed parameter files for each search are provided in the Supporting Information.

**Figure 2:**
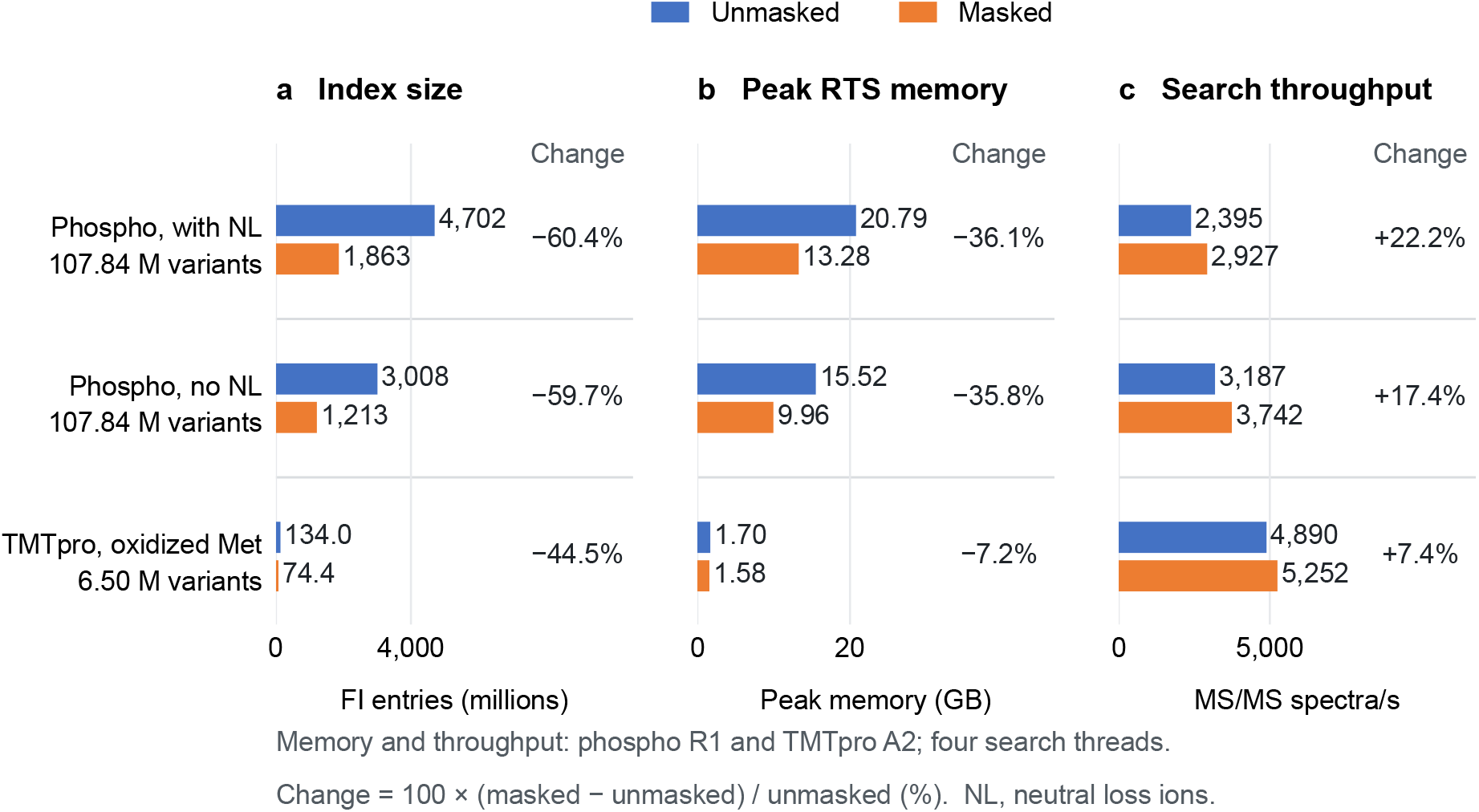
Effect of Carafe-informed masking on fragment ion index size, peak memory and search throughput. (a) Fragment ion (FI) index entries, (b) peak RTS process memory (GB) and (c) MS/MS search throughput (4 search threads) for unmasked (blue) versus masked (orange) FI searches. Peptide variant counts are shown below each search configuration; M denotes millions. Percentage changes are relative to the unmasked search. Memory and throughput are shown for the first acquisition of each dataset (phospho R1, TMTpro A2); the second acquisition of each dataset gave the same index and changes of similar size, with peak memory agreeing within 1.3 GB.

**Figure 3:**
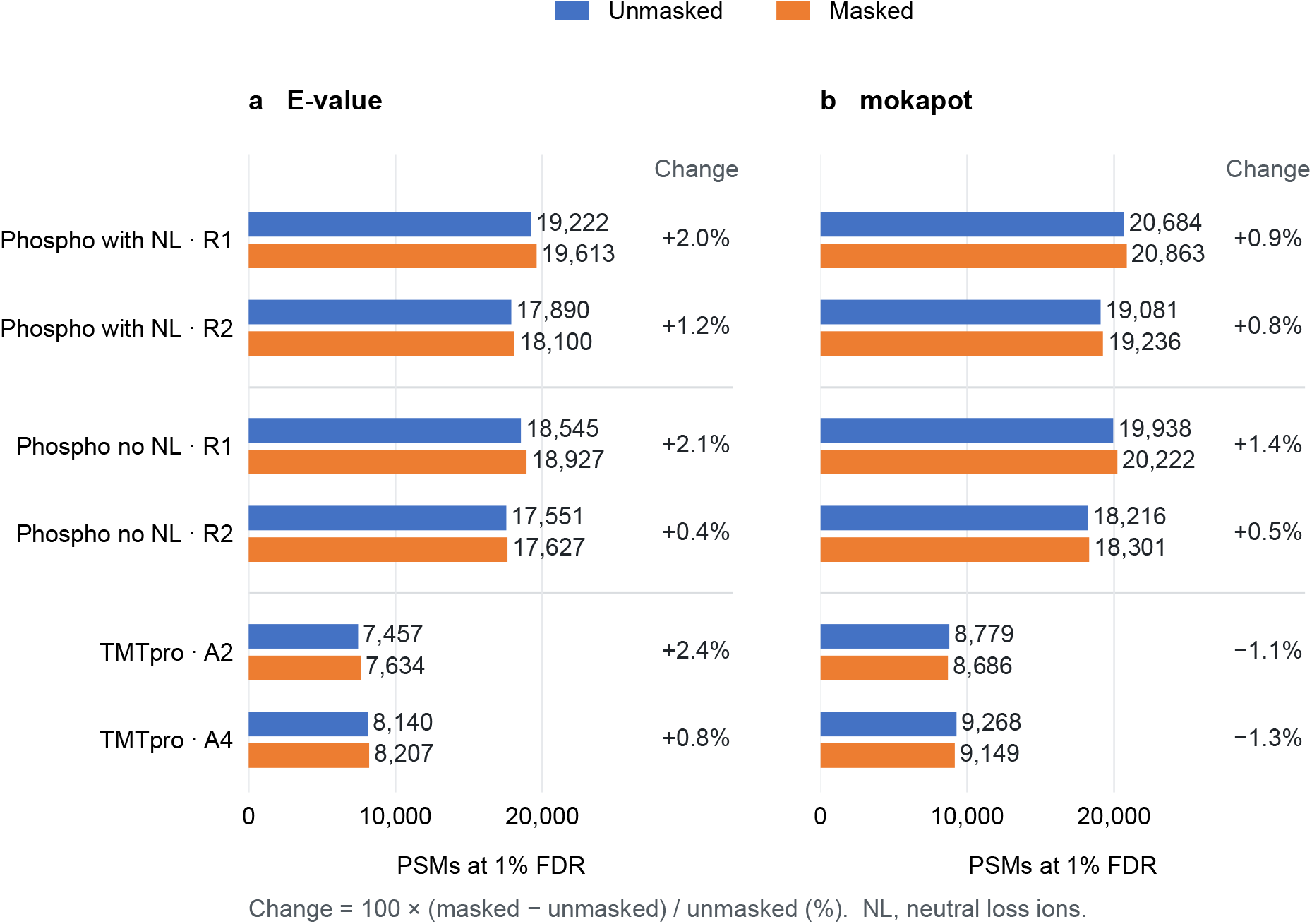
PSMs passing 1% FDR in unmasked versus masked FI searches. FDR was estimated by target-decoy competition on top-ranked PSMs, ranking by (a) Comet E-value or (b) mokapot rescoring. Blue and orange bars indicate unmasked and masked searches, respectively; percentage changes are relative to the unmasked search. R1 and R2, phosphoproteomic replicates, searched with (with NL) and without (no NL) the phosphate neutral loss ions indexed; A2 and A4, TMTpro fractions.

Carafe predictions were generated for precursor charges 2 and 3 of each peptidoform. The phosphorylation search space used the Carafe phosphorylation model with the instrument setting that matched the acquisition (Q Exactive, NCE 25). For the TMTpro search space, a CID-fragmentation TMT model supplied by Carafe was used (Lumos, NCE 35). The raw files were searched through Comet’s RTS framework using the C# real-time search application with four search threads. The application localizes the variable modifications of each top-ranked PSM with a C++ port of AScorePro ^21^ and reports the localized modification sites, while the XCorr reported is that of the positional isomer ranked first by the search. For each dataset, an unmasked and a masked fragment ion index search were run sequentially with the same plain peptide index and parameters, differing only in the presence of the mask.

False discovery rates were estimated by target-decoy competition on the top-ranked PSM of each spectrum, using two scores: Comet’s E-value and a mokapot ^22^ rescoring. A PSM was classified as a decoy only if every protein it maps to is a decoy sequence; hence, a peptide that is present in both a target and a decoy protein is considered a target. For E-value ranking, PSMs were sorted by ascending E-value and the FDR at each position in the list was estimated as the number of decoy PSMs plus one, divided by the number of target PSMs at or above that position; the q-value of each PSM is the minimum FDR at its position or any lower position, and the reported count is the number of target PSMs with q-value *≤* 0.01. For mokapot rescoring, each result file was converted to an input file with the following features: XCorr, ln(E-value), the absolute precursor mass error in ppm after allowing ^13^C isotope offsets of 0, 1 and 2, peptide length, number of missed cleavages, number of variable modifications, and an indicator variable for each precursor charge state present in the file. mokapot (version 0.10.1.dev8) was run separately on each result file with its default linear support vector machine model, three-fold cross-validation, training FDR of 1%, automatic choice of the initial ranking feature (which selected ln(E-value) in every run), a test FDR of 1% and a fixed random seed, and the reported count is the number of target PSMs with a mokapot PSM-level q-value *≤* 0.01. mokapot therefore rescored the single top-ranked PSM of each spectrum and did not re-rank lower-scoring candidates, which the real-time search does not report. The same procedures were applied to the masked and unmasked results of every configuration. Wall-clock time and peak memory of every pipeline stage and search were recorded externally: wall clock from process start to exit, and peak memory was estimated as the maximum total working set across the complete process tree, sampled once per second; index sizes, initialization time, and MS/MS search throughput were taken from Comet’s runtime statistics. All stages ran natively on a Windows 11 desktop (Intel Core Ultra 7 265K CPU and 64 GB RAM) with no other load. Masked and unmasked searches were executed sequentially. Agreement between masked and unmasked searches was assessed by keying the PSMs passing a 1% FDR (E-value ranking, and separately mokapot q-value) by scan number and comparing the assigned peptidoforms.

## 2 Results and discussion

### 2.1 Masking reduces index size and peak memory

We first evaluate the impact of masking on the fragment ion index size and peak memory usage. For the fragment ion index, masking reduced the number of entries written to the in-memory index relative to the corresponding unmasked search: by 44.5% for the TMTpro search (1.34 *×* 10^8^ to 7.44 *×* 10^7^ entries), by 60.4% for the phosphoproteomic search with neutral loss ions (4.70 *×* 10^9^ to 1.86 *×* 10^9^ entries), and by 59.7% for the same phosphoproteomic search without neutral loss ions (3.01 *×* 10^9^ to 1.21 *×* 10^9^ entries) as reported in Figure 2. The reduction in the phosphoproteomic space was nearly identical with and without neutral loss ions, so the larger pruning there is not attributable to the neutral loss series itself. Whether the phosphoproteomic index is pruned more heavily than the TMTpro index because the predicted spectra of phosphorylated peptides are sparser, because the two search spaces use different prediction models, or because they use different fragmentation modes and bin widths cannot be separated with these two datasets. For the TMTpro search the index is small relative to the rest of the RTS process (the preloaded spectra and the .NET runtime), so peak process memory was already modest at 1.70 GB for fraction A2, and masking reduced it by 7% to 1.58 GB. For the phosphoproteomic search (replicate R1), masking reduced peak process memory by 36%, from 20.8 GB to 13.3 GB, a saving of 7.5 GB that brings this search space within the memory budget of an acquisition computer. Without the neutral loss ions, the same phosphoproteomic index is smaller, 15.5 GB unmasked, and masking reduced it by 36% to 10.0 GB. The second acquisition of each dataset gave slightly smaller reductions (33.3% and 32.0% for the phosphoproteomic replicates with and without neutral loss ions, 7.0% for TMTpro A4), with peak memory agreeing within 1.3 GB.

### 2.2 Masking increases search throughput

We next examine the impact of masking on search speed. As expected, a smaller index is faster to search, and masking increased MS/MS search throughput in every search: by 22.2% and 18.8% for the two phospho-proteomic replicates with neutral loss ions, by 17.4% and 13.1% without neutral loss ions, and by 7.4% and 6.5% for the two TMTpro fractions (Figure 2). These gains are smaller than the reductions in index size and are expected because index lookup accounts for only part of the per-spectrum search time. Per-spectrum search times, recorded by the RTS application for every spectrum with a reported match, remained in the millisecond range: in the phosphoproteomic searches with neutral loss ions the 99th percentile fell from 9 ms to 7 ms with masking (maximum 106 and 105 ms), without neutral loss ions it was 4 ms unmasked and 3–4 ms masked, and in the TMTpro fractions it was 1 ms in both cases; no spectrum in any search approached the application’s 500 ms per-spectrum timeout limit. Masked searches also incur a fixed cost for loading the mask and caching its per-variant decisions during index construction: about 8 s of RTS initialization for the 107.8 million entry phosphoproteomic mask (4.5 GB on disk) and 0.5 s for the 6.5 million entry TMTpro mask (273 MB).

### 2.3 Masking is not detrimental to detection performance

Despite removing large numbers of low-intensity fragment entries, masking left the number of PSMs passing a 1% FDR threshold essentially unchanged, within *−*1.3% to +2.4% of the unmasked count (Figure 3). Ranked by E-value, masked searches identified 2.0% and 1.2% more PSMs in the phosphoproteomic replicates with neutral loss ions, 2.1% and 0.4% more without neutral loss ions, and 2.4% and 0.8% more in the TMTpro fractions. With mokapot rescoring, the phosphoproteomic gains were 0.9% and 0.8% with neutral loss ions and 1.4% and 0.5% without, while the TMTpro counts were 1.1% and 1.3% below the unmasked values. The detections gained by masking were almost entirely spectra to which the unmasked search had assigned the same peptidoform at a score just below the threshold. Masking changes which candidates are scored, not how a candidate is scored. For scans for which both searches reported the same peptidoform, the XCorr was identical in 97.7% (phosphoproteomic R1) and 99.9% (TMTpro A2) of cases; the exceptions are scans for which the two searches ranked different site isomers first before AScorePro localization. The gain instead reflects fewer and lower-scoring decoy matches. Masking reduced the number of rank-1 decoy PSMs by 25% in the phosphoproteomic replicates (8,397 to 6,299 and 6,286 to 4,698) and shifted the decoy XCorr distribution downward (90th percentile 1.37 to 1.35), so that the same correct detections clear the FDR threshold. Figure 4 shows the paired XCorr values and the target and decoy score distributions.

**Figure 4:**
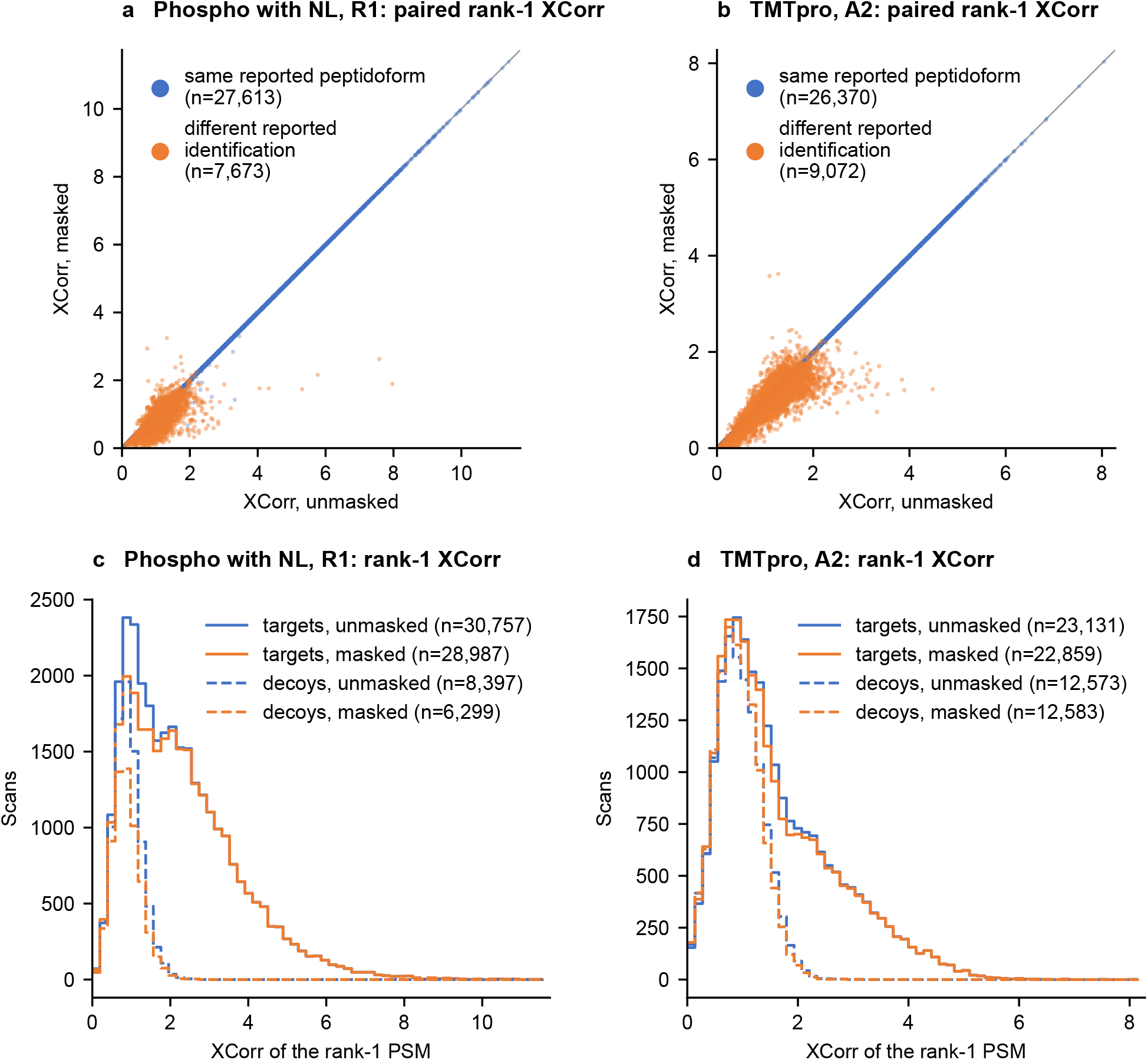
XCorr before and after masking. (a, b) Rank-1 XCorr of every scan with a PSM in both the unmasked and the masked search, for phospho with NL R1 (a) and TMTpro A2 (b). Scans for which both searches reported the same peptidoform (blue) lie on the diagonal because masking does not alter the score of a candidate; the few blue points off the diagonal are scans for which the searches ranked different site isomers first before AScorePro localization. Scans for which the reported detection differs (orange), in most cases a different peptide and in a few cases different modification sites on the same peptide, are confined to low XCorr and tend to score lower after masking. A further 3,868 (a) and 262 (b) scans had a rank-1 PSM only in the unmasked search, with median XCorr 0.61 and 0.20. (c, d) Distributions of rank-1 XCorr for target (solid) and decoy (dashed) PSMs in the unmasked (blue) and masked (orange) searches. Masking removes low-scoring candidates and shrinks the high-scoring tail of the decoy distribution (and, in the phosphoproteomic search, the decoy population itself) while leaving confident target scores unchanged.

### 2.4 Masked and unmasked searches agree on the vast majority of detections

A comparison of PSMs passing 1% FDR (E-value ranking) showed very high agreement between masked and unmasked searches. Among scans confidently identified by both, peptide assignments were identical for 99.94% and 99.97% of shared scans in the two phosphoproteomic replicates with neutral loss ions (12 and 6 disagreements among 18,843 and 17,529 shared scans), for 99.86% and 99.78% without neutral loss ions (26 and 38 disagreements among 18,167 and 17,143), and for 100% and 99.99% in the TMTpro fractions (0 and 1 disagreements among 7,306 and 7,943). Of the phosphoproteomic PSMs found only by the masked search, 95–97% (747 of 770 in R1 and 555 of 571 in R2 with neutral loss ions; 735 of 760 and 460 of 484 without) had received the same peptidoform from the unmasked search at a score below the threshold. Agreement was equally high when PSMs were selected by mokapot q-value (99.8–99.96% identical assignments among 17,787–20,441 shared scans in the phosphoproteomic searches and 99.98% among 8,568 and 9,048 in the TMTpro fractions).

### 2.5 Neutral loss aware masking extends Comet’s fragment ion index support

The masking framework also extends Comet’s fragment ion index, which previously contained only conventional fragment ion series, to modification-specific neutral loss fragments. Carafe predicts the intensities of both unshifted and neutral loss shifted b-ions and y-ions, including the dominant phosphopeptide neutral loss species, so these ions can be queried through the same mechanism and in the same index as the base ions. To evaluate what the neutral loss ions cost and what they contribute, the phosphoproteomic search space was indexed with and without the neutral loss channel enabled. Indexing the neutral loss ions enlarged the unmasked index by 56% (3.01 *×* 10^9^ to 4.70 *×* 10^9^) and peak memory by 34% (15.5 GB to 20.8 GB), lowered search throughput by 24–25%, and increased the number of PSMs accepted at 1% FDR by 3.7% and 1.9% (E-value) and by 3.7% and 4.7% (mokapot) (Figures 2 and 3). From a reduction perspective, masking acted almost identically in both configurations with index reductions of 60.4% and 59.7%, and memory saving of 36% in both. This shows that masking is as effective for neutral loss fragments as for the base ion series. In fact, the masked index with neutral loss ions was smaller than the unmasked index without them (1.86 *×* 10^9^ versus 3.01 *×* 10^9^ entries; 13.3 versus 15.5 GB) while identifying 5.8% and 3.1% more PSMs by E-value and 4.6% and 5.6% more by mokapot, at a search rate 8–10% lower.

### 2.6 One time inference costs are amortized across searches

Carafe inference is the most computationally intensive step of this analysis procedure. On an Intel Core Ultra 7 265K processor, running three concurrent 8-thread Carafe processes required 2.1 h for the TMTpro search space (6.5 million variants, 13.0 million peptidoform-charge rows) and 59.6 h for the phosphoproteomic search space (107.8 million variants, 215.7 million peptidoform-charge rows). GPU inference would be substantially faster. Every other pipeline step took minutes: for the phosphoproteomic space, plain index construction, variant export and format conversion together took 23.3 min, prediction store generation 13.3 min and mask construction 6.7 min. A second mask for the same search space without neutral loss ions reused the prediction store and needed 23 min of index preparation and 6 min of mask construction. For the TMTpro space, the times were 1.3 min, 34 s, and 19 s respectively. This cost is incurred once per protein database and modification configuration, and the mask can be reused for any dataset searched with the same database, modification configuration and index settings, which the mask header validates at load time. Machine learning inference is thereby fully detached from the time-critical search path. The only search-time overhead is loading and consulting the precomputed mask, which added about 8 s (phosphoproteomic, 4.5 GB mask) and 0.5 s (TMTpro, 273 MB mask) to RTS initialization.

## 3 Conclusions

Carafe-guided fragment ion masking uses deep learning predictions of fragment ion intensities, including neutral loss species, to prune Comet’s fragment ion index before database searching. In the search configurations evaluated, it reduced the in-memory index by 45–60% and peak search memory by 7–36%, making large search spaces more practical and concurrent fragment ion index searches during acquisition feasible. Masking also increased MS/MS search throughput by 6–22%, kept per-spectrum search times below 10 ms at the 99th percentile, and left the number of confident PSMs at 1% FDR essentially unchanged (*−*1% to +2% across datasets and scoring methods). Because all inference is performed in a one-time, ahead-of-time pipeline, its cost is amortized across subsequent searches of the same database and modification configuration, adding no per-spectrum overhead and only a few seconds of startup latency, so the approach meets the per-spectrum time requirements of RTS.

## Data and software availability

Comet-FI and Carafe are open source under the Apache License Version 2.0 and are available at https://github.com/UWPR/Comet and https://github.com/Noble-Lab/Carafe. Searches used Comet built from the carafe branch of the Comet repository at commit 29096659, which also contains the mask-generation tools; Carafe predictions used the `tmt` branch of the Carafe repository at commit a9c5049. The MS/MS data files were downloaded from PRIDE with the accession numbers PXD043660 ^18^ and PXD011175 ^19^.

## Acknowledgments

This work was supported by the National Science Foundation award 2245300 (W.S.N.), the National Science Foundation Graduate Research Fellowship Program (Grant No. DGE-2140004, B.W.), and the National Institutes of Health grant R24GM141156. D.K.S. acknowledges support from the NIH/NIGMS (R35GM150919), Washington Research Foundation, the W. M. Keck Foundation, an Andy Hill CARE Distinguished Researcher Award, a Cancer Consortium New Investigator Award, and the Pew Charitable Trusts.

This work is supported in part by the University of Washington’s Proteomics Resource (UWPR95794). Software development, including both the C++ integration and the accompanying Python scripts, was assisted by Claude Code (Anthropic); all code contributions were subject to author oversight, review, testing, and validation.

## Author contributions

|  | JKE | MRH | DKS | MJM | WSN | BW |
|---|---|---|---|---|---|---|
| Conceptualization | ● | ● |  |  |  | ● |
| Data curation | ● |  | ● |  |  | ● |
| Funding acquisition |  |  | ● | ● | ● | ● |
| Investigation | ● |  |  |  | ● | ● |
| Methodology | ● |  |  |  |  | ● |
| Resources | ● |  |  | ● | ● | ● |
| Software | ● |  |  |  |  | ● |
| Supervision | ● |  | ● | ● | ● | ● |
| Validation | ● |  |  |  |  | ● |
| Visualization | ● |  |  |  |  | ● |
| Writing—original draft | ● |  |  |  |  | ● |
| Writing—review & editing | ● | ● | ● | ● | ● | ● |

## Conflict of interest

The MacCoss Lab at the University of Washington receives funding from Agilent, Bruker, Sciex, Shimadzu, Thermo Fisher Scientific, and Waters to support the development of Skyline, a quantitative analysis software tool. M.J.M. is a paid consultant for Thermo Fisher Scientific. D.K.S. is a collaborator and/or consultant for Thermo Fisher Scientific, Genentech, and AI Proteins.

## Notes

https://github.com/UWPR/Comet

