## Supporting Information for "Improving real-time search through deep learning-enhanced fragment ion indexing"

### Supporting Information for:
### Improving real-time search through deep learning-enhanced fragment ion indexing

#### S1. Search configurations

Three search configurations were evaluated: the phosphoproteomic search space with the phosphate neutral loss ions indexed ("with NL"), the same search space without them ("no NL"), and the TMTpro search space. Each configuration is fully specified by one Comet parameter file, reproduced verbatim in Sections S2–S4. The three files share the same template; Table S1 lists the parameters that differ.

**Table S1.** Parameters that differ between the three configurations.

| Parameter | Phospho, with NL | Phospho, no NL | TMTpro |
|---|---|---|---|
| `peptide_length_range` | 7 40 | 7 40 | 7 50 |
| `variable_mod02` (STY phospho) | 79.966331 STY, max 3, NL 97.976896 | 79.966331 STY, max 3, NL 0.0 | none |
| `add_Nterm_peptide` | 0.0 | 0.0 | 304.207146 (TMTpro) |
| `add_K_lysine` | 0.0 | 0.0 | 304.207146 (TMTpro) |
| `fragment_bin_tol` | 0.02 | 0.02 | 1.0005 |
| `fragment_bin_offset` | 0.0 | 0.0 | 0.4 |
| `theoretical_fragment_ions` | 0 (flanking peaks) | 0 (flanking peaks) | 1 (monoisotopic peak only) |

All three configurations allow methionine oxidation (+15.9949 Da; maximum 3 per peptide).

**Protein database.** All searches used `human.canonical.target-decoy.fasta`: the UniProt human canonical proteome (20,406 entries) plus 48 common contaminant proteins (bovine and porcine trypsin, Lys-C, chymotrypsinogens, pepsins, bovine serum albumin, caseins, sheep wool keratins, and others), for 20,454 target sequences, each followed by its reversed sequence with the `DECOY_` prefix (20,454 decoys; `decoy_search = 0` because the decoys are part of the database). Contaminant matches are counted as targets in every reported PSM count.

**How the parameter files were used.** Each file was used by `comet.exe -i` to build the plain peptide index (`.idx`) for its configuration and by the Carafe pipeline (`tools/carafe.py prerun`) to enumerate the peptidoform variants for prediction. The mask thresholds are the two `carafe_mask_*` keys near the end of each file (keep a predicted fragment at ≥10% of the variant's predicted base peak; always keep at least the 6 most intense). `fragment_index_predicted_mask_file` is left empty in the files because the mask was supplied on the command line of the real-time search application instead (see below), so that masked and unmasked searches used one identical index.

**Real-time searches.** The searches were run with the Comet project's C# `RealtimeSearch` application against the `.idx` of each configuration. The application reads the enzyme, modification, peptide length and mass settings from the index header and takes the remaining scoring settings from its command line, which was set to match the parameter file: `-threads 4 -isotope-error 2` for every run, plus `-fragment-bin-tol 1.0005 -fragment-bin-offset 0.4 -theoretical-fragment-ions 1` for the TMTpro ion-trap data, and `-mask <file>.fi_mask` for the masked run of each pair. Precursor tolerance (20 ppm), minimum peaks (10), maximum fragment charge (3), precursor charge range (1–6) and b/y ions are the same in the application and the parameter files.

**Carafe predictions.** Precursor charges 2 and 3 were predicted for every variant. Phosphoproteomic search space: Carafe phosphorylation mode (neutral loss channels on), instrument `QE`, NCE 25; the no-NL mask was built from the same predictions with the neutral loss channels ignored. TMTpro search space: the Carafe CID-fragmentation TMT model, instrument `Lumos`, NCE 35, general mode.

### S2. Phosphoproteomic search, with neutral loss ions (`comet.params`)

```
# comet_version 2026.01 rev. 1
# Comet MS/MS search engine parameters file.
# Everything following the '#' symbol is treated as a comment.
#
database_name = C:\Work\fasta\human.canonical.target-decoy.fasta
decoy_search = 0                       # 0=no (default), 1=internal decoy concatenated, 2=internal decoy separate

num_threads = 8                        # 0=poll CPU to set num threads; else specify num threads directly (max 128)

#
# fragment ion index; limited to 5 variable mods and up to 5 modified residues per mod
#
fragindex_min_ions_score = 3           # minimum number of matched fragment ion index peaks for scoring
fragindex_min_ions_report = 3          # minimum number of matched fragment ion index peaks for reporting(>=
↪ fragindex_min_ions_score)
fragindex_num_spectrumpeaks = 150      # number of peaks from spectrum to use for fragment ion index matching
fragindex_min_fragmentmass = 200.0     # low mass cutoff for fragment ions
fragindex_max_fragmentmass = 2000.0    # high mass cutoff for fragment ions
fragindex_skipreadprecursors = 1       # 0=read precursors to limit fragment ion index, 1=skip reading precursors (default)

#
# masses
#
peptide_mass_tolerance_upper = 20.0    # upper bound of the precursor mass tolerance
peptide_mass_tolerance_lower = -20.0   # lower bound of the precursor mass tolerance; USUALLY NEGATIVE TO BE LOWER THAN 0
peptide_mass_units = 2                 # 0=amu, 1=mmu, 2=ppm
precursor_tolerance_type = 1           # 0=MH+ (default), 1=precursor m/z; only valid for amu/mmu tolerances
isotope_error = 2                      # 0=off, 1=0/1 (C13 error), 2=0/1/2, 3=0/1/2/3, 4=-1/0/1/2/3, 5=-1/0/1

#
# search enzyme
#
search_enzyme_number = 1               # choose from list at end of this params file
search_enzyme2_number = 0              # second enzyme; set to 0 if no second enzyme
sample_enzyme_number = 1               # specifies the sample enzyme which is possibly different than the one applied to the
↪ search;
                                       # used by PeptideProphet to calculate NTT & NMC in pepXML output (default=1 for
↪ trypsin).
num_enzyme_termini = 2                 # 1 (semi-digested), 2 (fully digested, default), 8 C-term unspecific , 9 N-term
↪ unspecific
allowed_missed_cleavage = 2            # maximum value is 5; for enzyme search

#
# Up to 15 variable_mod entries are supported for a standard search; manually add additional entries as needed
# format:  <mass> <residues> <0=variable/else binary> <max_mods_per_peptide> <term_distance> <n/c-term> <required>
↪ <neutral_loss>
#     e.g. 79.966331 STY 0 3 -1 0 0 97.976896
#
variable_mod01 = 15.9949 M 0 3 -1 0 0 0.0
variable_mod02 = 79.966331 STY 0 3 -1 0 0 97.976896
variable_mod03 = 0.0 X 0 3 -1 0 0 0.0
variable_mod04 = 0.0 X 0 3 -1 0 0 0.0
variable_mod05 = 0.0 X 0 3 -1 0 0 0.0
max_variable_mods_in_peptide = 3
require_variable_mod = 0

#
# fragment ions
#
# ion trap ms/ms:  1.0005 tolerance, 0.4 offset (mono masses), theoretical_fragment_ions = 1
# high res ms/ms:    0.02 tolerance, 0.0 offset (mono masses), theoretical_fragment_ions = 0, spectrum_batch_size = 15000
#
fragment_bin_tol = 0.02                # binning to use on fragment ions
fragment_bin_offset = 0.0              # offset position to start the binning (0.0 to 1.0)
theoretical_fragment_ions = 0          # 0=use flanking peaks, 1=M peak only
use_A_ions = 0
use_B_ions = 1
use_C_ions = 0
use_X_ions = 0
use_Y_ions = 1
use_Z_ions = 0
use_Z1_ions = 0
use_NL_ions = 0                        # 0=no, 1=yes to consider NH3/H2O neutral loss peaks

#
# output
#
```

```
output_sqtfile = 0                      # 0=no, 1=yes  write sqt file
output_txtfile = 1                      # 0=no, 1=yes, 2=Crux-formatted  write tab-delimited txt file
output_pepxmlfile = 0                   # 0=no, 1=yes  write pepXML file
output_mzidentmlfile = 0                # 0=no, 1=yes  write mzIdentML file
output_percolatorfile = 0               # 0=no, 1=yes  write Percolator pin file
num_output_lines = 1                    # num peptide results to show

#
# mzXML/mzML/raw file parameters
#
scan_range = 0 0                        # start and end scan range to search; either entry can be set independently
precursor_charge = 0 0                  # precursor charge range to analyze; does not override any existing charge; 0 as 1st
↪ entry ignores parameter
override_charge = 0                     # 0=no, 1=override precursor charge states, 2=ignore precursor charges outside
↪ precursor_charge range, 3=see online
ms_level = 2                            # MS level to analyze, valid are levels 2 (default) or 3
activation_method = ALL                 # activation method; used if activation method set; allowed ALL, CID, ECD, ETD,
↪ ETD+SA, PQD, HCD, IRMPD, SID

#
# misc parameters
#
digest_mass_range = 700.0 5000.0        # MH+ peptide mass range to analyze
peptide_length_range = 7 40             # minimum and maximum peptide length to analyze (default min 1 to allowed max 51)
max_duplicate_proteins = 10             # maximum number of additional duplicate protein names to report for each peptide ID;
↪ -1 reports all duplicates
max_fragment_charge = 3                 # set maximum fragment charge state to analyze (allowed max 5)
min_precursor_charge = 1                # set minimum precursor charge state to analyze (1 if missing)
max_precursor_charge = 6                # set maximum precursor charge state to analyze (allowed max 9)
clip_nterm_methionine = 0               # 0=leave protein sequences as-is; 1=also consider sequence w/o N-term methionine
spectrum_batch_size = 75000             # max. # of spectra to search at a time; 0 to search the entire scan range in one loop
decoy_prefix = DECOY_                   # decoy entries are denoted by this string which is pre-pended to each protein
↪ accession
equal_I_and_L = 1                       # 0=treat I and L as different; 1=treat I and L as same
mass_offsets =                          # one or more mass offsets to search (values substracted from deconvoluted precursor
↪ mass)

#
# spectral processing
#
minimum_peaks = 10                      # required minimum number of peaks in spectrum to search (default 10)
minimum_intensity = 0                  # minimum intensity value to read in
remove_precursor_peak = 0               # 0=no, 1=yes, 2=all charge reduced precursor peaks (for ETD), 3=phosphate neutral
↪ loss peaks
remove_precursor_tolerance = 1.5        # +- Da tolerance for precursor removal
clear_mz_range = 0.0 0.0                # clear out all peaks in the specified m/z range e.g. remove reporter ion region of
↪ TMT spectra
percentage_base_peak = 0.0              # specify a percentage (e.g. "0.05" for 5%) of the base peak intensity as a minimum
↪ intensity threshold

#
# static modifications
#
add_Cterm_peptide = 0.0
add_Nterm_peptide = 0.0
add_Cterm_protein = 0.0
add_Nterm_protein = 0.0

add_G_glycine = 0.0000                  # added to G - avg.  57.0513, mono.  57.02146
add_A_alanine = 0.0000                  # added to A - avg.  71.0779, mono.  71.03711
add_S_serine = 0.0000                   # added to S - avg.  87.0773, mono.  87.03203
add_P_proline = 0.0000                  # added to P - avg.  97.1152, mono.  97.05276
add_V_valine = 0.0000                   # added to V - avg.  99.1311, mono.  99.06841
add_T_threonine = 0.0000                # added to T - avg. 101.1038, mono. 101.04768
add_C_cysteine = 57.021464              # added to C - avg. 103.1429, mono. 103.00918
add_L_leucine = 0.0000                  # added to L - avg. 113.1576, mono. 113.08406
add_I_isoleucine = 0.0000               # added to I - avg. 113.1576, mono. 113.08406
add_N_asparagine = 0.0000               # added to N - avg. 114.1026, mono. 114.04293
add_D_aspartic_acid = 0.0000            # added to D - avg. 115.0874, mono. 115.02694
add_Q_glutamine = 0.0000                # added to Q - avg. 128.1292, mono. 128.05858
add_K_lysine = 0.0000                   # added to K - avg. 128.1723, mono. 128.09496
add_E_glutamic_acid = 0.0000            # added to E - avg. 129.1140, mono. 129.04259
add_M_methionine = 0.0000               # added to M - avg. 131.1961, mono. 131.04048
add_H_histidine = 0.0000                # added to H - avg. 137.1393, mono. 137.05891
add_F_phenylalanine = 0.0000            # added to F - avg. 147.1739, mono. 147.06841
add_U_selenocysteine = 0.0000           # added to U - avg. 150.0379, mono. 150.95363
add_R_arginine = 0.0000                 # added to R - avg. 156.1857, mono. 156.10111
add_Y_tyrosine = 0.0000                 # added to Y - avg. 163.0633, mono. 163.06333
add_W_tryptophan = 0.0000               # added to W - avg. 186.0793, mono. 186.07931
add_O_pyrrolysine = 0.0000              # added to O - avg. 237.2982, mono  237.14773
add_B_user_amino_acid = 0.0000          # added to B - avg.   0.0000, mono.   0.00000
```

```
add_J_user_amino_acid = 0.0000         # added to J - avg.   0.0000, mono.   0.00000
add_X_user_amino_acid = 0.0000         # added to X - avg.   0.0000, mono.   0.00000
add_Z_user_amino_acid = 0.0000         # added to Z - avg.   0.0000, mono.   0.00000

fragment_index_predicted_mask_file =   # Carafe predicted-fragment mask (.fi_mask); empty = disabled. Set by run_rts.ps1 via
↪ RealtimeSearch.exe --mask; kept empty here for the .idx build
carafe_mask_min_relative_intensity = 0.10   # read by tools/carafe.py prerun --params only
carafe_mask_min_peaks = 6                   # read by tools/carafe.py prerun --params only
#
# COMET_ENZYME_INFO _must_ be at the end of this parameters file
# Enzyme entries can be added/deleted/edited

#
[COMET_ENZYME_INFO]
0.  Cut_everywhere         0      -           -
1.  Trypsin                1      KR          P
2.  Trypsin/P              1      KR          -
3.  Lys_C                  1      K           P
4.  Lys_N                  0      K           -
5.  Arg_C                  1      R           P
6.  Asp_N                  0      DN          -
7.  CNBr                   1      M           -
8.  Asp-N_ambic            1      DE          -
9.  PepsinA                1      FL          -
10. Chymotrypsin           1      FWYL        P
11. No_cut                 1      @           @
```

### S3. Phosphoproteomic search, no neutral loss ions (`comet.params.nonl`)

Identical to S2 except that the neutral loss of `variable_mod02` is 0.0.

```
# comet_version 2026.01 rev. 1
# Comet MS/MS search engine parameters file.
# Everything following the '#' symbol is treated as a comment.
#
database_name = C:\Work\fasta\human.canonical.target-decoy.fasta
decoy_search = 0                       # 0=no (default), 1=internal decoy concatenated, 2=internal decoy separate

num_threads = 8                        # 0=poll CPU to set num threads; else specify num threads directly (max 128)

#
# fragment ion index; limited to 5 variable mods and up to 5 modified residues per mod
#
fragindex_min_ions_score = 3           # minimum number of matched fragment ion index peaks for scoring
fragindex_min_ions_report = 3          # minimum number of matched fragment ion index peaks for reporting(>=
↪ fragindex_min_ions_score)
fragindex_num_spectrumpeaks = 150      # number of peaks from spectrum to use for fragment ion index matching
fragindex_min_fragmentmass = 200.0     # low mass cutoff for fragment ions
fragindex_max_fragmentmass = 2000.0    # high mass cutoff for fragment ions
fragindex_skipreadprecursors = 1       # 0=read precursors to limit fragment ion index, 1=skip reading precursors (default)

#
# masses
#
peptide_mass_tolerance_upper = 20.0    # upper bound of the precursor mass tolerance
peptide_mass_tolerance_lower = -20.0   # lower bound of the precursor mass tolerance; USUALLY NEGATIVE TO BE LOWER THAN 0
peptide_mass_units = 2                 # 0=amu, 1=mmu, 2=ppm
precursor_tolerance_type = 1           # 0=MH+ (default), 1=precursor m/z; only valid for amu/mmu tolerances
isotope_error = 2                      # 0=off, 1=0/1 (C13 error), 2=0/1/2, 3=0/1/2/3, 4=-1/0/1/2/3, 5=-1/0/1

#
# search enzyme
#
search_enzyme_number = 1               # choose from list at end of this params file
search_enzyme2_number = 0              # second enzyme; set to 0 if no second enzyme
sample_enzyme_number = 1               # specifies the sample enzyme which is possibly different than the one applied to the
↪ search;
                                       # used by PeptideProphet to calculate NTT & NMC in pepXML output (default=1 for
↪ trypsin).
num_enzyme_termini = 2                 # 1 (semi-digested), 2 (fully digested, default), 8 C-term unspecific , 9 N-term
↪ unspecific
allowed_missed_cleavage = 2            # maximum value is 5; for enzyme search

#
# Up to 15 variable_mod entries are supported for a standard search; manually add additional entries as needed
# format:  <mass> <residues> <0=variable/else binary> <max_mods_per_peptide> <term_distance> <n/c-term> <required>
↪ <neutral_loss>
#     e.g. 79.966331 STY 0 3 -1 0 0 97.976896
#
variable_mod01 = 15.9949 M 0 3 -1 0 0 0.0
variable_mod02 = 79.966331 STY 0 3 -1 0 0 0.0
variable_mod03 = 0.0 X 0 3 -1 0 0 0.0
variable_mod04 = 0.0 X 0 3 -1 0 0 0.0
variable_mod05 = 0.0 X 0 3 -1 0 0 0.0
max_variable_mods_in_peptide = 3
require_variable_mod = 0

#
# fragment ions
#
# ion trap ms/ms:  1.0005 tolerance, 0.4 offset (mono masses), theoretical_fragment_ions = 1
# high res ms/ms:    0.02 tolerance, 0.0 offset (mono masses), theoretical_fragment_ions = 0, spectrum_batch_size = 15000
#
fragment_bin_tol = 0.02                # binning to use on fragment ions
fragment_bin_offset = 0.0              # offset position to start the binning (0.0 to 1.0)
theoretical_fragment_ions = 0          # 0=use flanking peaks, 1=M peak only
use_A_ions = 0
use_B_ions = 1
use_C_ions = 0
use_X_ions = 0
use_Y_ions = 1
use_Z_ions = 0
use_Z1_ions = 0
use_NL_ions = 0                        # 0=no, 1=yes to consider NH3/H2O neutral loss peaks

#
# output
```

```
#
output_sqtfile = 0                       # 0=no, 1=yes  write sqt file
output_txtfile = 1                       # 0=no, 1=yes, 2=Crux-formatted  write tab-delimited txt file
output_pepxmlfile = 0                    # 0=no, 1=yes  write pepXML file
output_mzidentmlfile = 0                 # 0=no, 1=yes  write mzIdentML file
output_percolatorfile = 0                # 0=no, 1=yes  write Percolator pin file
num_output_lines = 1                     # num peptide results to show

#
# mzXML/mzML/raw file parameters
#
scan_range = 0 0                         # start and end scan range to search; either entry can be set independently
precursor_charge = 0 0                   # precursor charge range to analyze; does not override any existing charge; 0 as 1st
↪ entry ignores parameter
override_charge = 0                      # 0=no, 1=override precursor charge states, 2=ignore precursor charges outside
↪ precursor_charge range, 3=see online
ms_level = 2                             # MS level to analyze, valid are levels 2 (default) or 3
activation_method = ALL                  # activation method; used if activation method set; allowed ALL, CID, ECD, ETD,
↪ ETD+SA, PQD, HCD, IRMPD, SID

#
# misc parameters
#
digest_mass_range = 700.0 5000.0         # MH+ peptide mass range to analyze
peptide_length_range = 7 40              # minimum and maximum peptide length to analyze (default min 1 to allowed max 51)
max_duplicate_proteins = 10              # maximum number of additional duplicate protein names to report for each peptide ID;
↪ -1 reports all duplicates
max_fragment_charge = 3                  # set maximum fragment charge state to analyze (allowed max 5)
min_precursor_charge = 1                 # set minimum precursor charge state to analyze (1 if missing)
max_precursor_charge = 6                 # set maximum precursor charge state to analyze (allowed max 9)
clip_nterm_methionine = 0                # 0=leave protein sequences as-is; 1=also consider sequence w/o N-term methionine
spectrum_batch_size = 75000              # max. # of spectra to search at a time; 0 to search the entire scan range in one loop
decoy_prefix = DECOY_                    # decoy entries are denoted by this string which is pre-pended to each protein
↪ accession
equal_I_and_L = 1                        # 0=treat I and L as different; 1=treat I and L as same
mass_offsets =                           # one or more mass offsets to search (values substracted from deconvoluted precursor
↪ mass)

#
# spectral processing
#
minimum_peaks = 10                       # required minimum number of peaks in spectrum to search (default 10)
minimum_intensity = 0                   # minimum intensity value to read in
remove_precursor_peak = 0                # 0=no, 1=yes, 2=all charge reduced precursor peaks (for ETD), 3=phosphate neutral
↪ loss peaks
remove_precursor_tolerance = 1.5         # +- Da tolerance for precursor removal
clear_mz_range = 0.0 0.0                 # clear out all peaks in the specified m/z range e.g. remove reporter ion region of
↪ TMT spectra
percentage_base_peak = 0.0               # specify a percentage (e.g. "0.05" for 5%) of the base peak intensity as a minimum
↪ intensity threshold

#
# static modifications
#
add_Cterm_peptide = 0.0
add_Nterm_peptide = 0.0
add_Cterm_protein = 0.0
add_Nterm_protein = 0.0

add_G_glycine = 0.0000                   # added to G - avg.  57.0513, mono.  57.02146
add_A_alanine = 0.0000                   # added to A - avg.  71.0779, mono.  71.03711
add_S_serine = 0.0000                    # added to S - avg.  87.0773, mono.  87.03203
add_P_proline = 0.0000                   # added to P - avg.  97.1152, mono.  97.05276
add_V_valine = 0.0000                    # added to V - avg.  99.1311, mono.  99.06841
add_T_threonine = 0.0000                 # added to T - avg. 101.1038, mono. 101.04768
add_C_cysteine = 57.021464               # added to C - avg. 103.1429, mono. 103.00918
add_L_leucine = 0.0000                   # added to L - avg. 113.1576, mono. 113.08406
add_I_isoleucine = 0.0000                # added to I - avg. 113.1576, mono. 113.08406
add_N_asparagine = 0.0000                # added to N - avg. 114.1026, mono. 114.04293
add_D_aspartic_acid = 0.0000             # added to D - avg. 115.0874, mono. 115.02694
add_Q_glutamine = 0.0000                 # added to Q - avg. 128.1292, mono. 128.05858
add_K_lysine = 0.0000                    # added to K - avg. 128.1723, mono. 128.09496
add_E_glutamic_acid = 0.0000             # added to E - avg. 129.1140, mono. 129.04259
add_M_methionine = 0.0000                # added to M - avg. 131.1961, mono. 131.04048
add_H_histidine = 0.0000                 # added to H - avg. 137.1393, mono. 137.05891
add_F_phenylalanine = 0.0000             # added to F - avg. 147.1739, mono. 147.06841
add_U_selenocysteine = 0.0000            # added to U - avg. 150.0379, mono. 150.95363
add_R_arginine = 0.0000                  # added to R - avg. 156.1857, mono. 156.10111
add_Y_tyrosine = 0.0000                  # added to Y - avg. 163.0633, mono. 163.06333
add_W_tryptophan = 0.0000                # added to W - avg. 186.0793, mono. 186.07931
add_O_pyrrolysine = 0.0000               # added to O - avg. 237.2982, mono  237.14773
```

```
add_B_user_amino_acid = 0.0000         # added to B - avg.   0.0000, mono.   0.00000
add_J_user_amino_acid = 0.0000         # added to J - avg.   0.0000, mono.   0.00000
add_X_user_amino_acid = 0.0000         # added to X - avg.   0.0000, mono.   0.00000
add_Z_user_amino_acid = 0.0000         # added to Z - avg.   0.0000, mono.   0.00000

fragment_index_predicted_mask_file =   # Carafe predicted-fragment mask (.fi_mask); empty = disabled. Set by run_rts.ps1 via
↪ RealtimeSearch.exe --mask; kept empty here for the .idx build
carafe_mask_min_relative_intensity = 0.10   # read by tools/carafe.py prerun --params only
carafe_mask_min_peaks = 6                   # read by tools/carafe.py prerun --params only
#
# COMET_ENZYME_INFO _must_ be at the end of this parameters file
# Enzyme entries can be added/deleted/edited

#
[COMET_ENZYME_INFO]
0.  Cut_everywhere         0      -           -
1.  Trypsin                1      KR          P
2.  Trypsin/P              1      KR          -
3.  Lys_C                  1      K           P
4.  Lys_N                  0      K           -
5.  Arg_C                  1      R           P
6.  Asp_N                  0      DN          -
7.  CNBr                   1      M           -
8.  Asp-N_ambic            1      DE          -
9.  PepsinA                1      FL          -
10. Chymotrypsin           1      FWYL        P
11. No_cut                 1      @           @
```

### S4. TMTpro search (comet.params)

```
# comet_version 2026.01 rev. 1
# Comet MS/MS search engine parameters file.
# Everything following the '#' symbol is treated as a comment.
#
database_name = C:\Work\fasta\human.canonical.target-decoy.fasta
decoy_search = 0                       # 0=no (default), 1=internal decoy concatenated, 2=internal decoy separate

num_threads = 8                        # 0=poll CPU to set num threads; else specify num threads directly (max 128)

#
# fragment ion index; limited to 5 variable mods and up to 5 modified residues per mod
#
fragindex_min_ions_score = 3           # minimum number of matched fragment ion index peaks for scoring
fragindex_min_ions_report = 3          # minimum number of matched fragment ion index peaks for reporting(>=
↪ fragindex_min_ions_score)
fragindex_num_spectrumpeaks = 150      # number of peaks from spectrum to use for fragment ion index matching
fragindex_min_fragmentmass = 200.0     # low mass cutoff for fragment ions
fragindex_max_fragmentmass = 2000.0    # high mass cutoff for fragment ions
fragindex_skipreadprecursors = 1       # 0=read precursors to limit fragment ion index, 1=skip reading precursors (default)

#
# masses
#
peptide_mass_tolerance_upper = 20.0    # upper bound of the precursor mass tolerance
peptide_mass_tolerance_lower = -20.0   # lower bound of the precursor mass tolerance; USUALLY NEGATIVE TO BE LOWER THAN 0
peptide_mass_units = 2                 # 0=amu, 1=mmu, 2=ppm
precursor_tolerance_type = 1           # 0=MH+ (default), 1=precursor m/z; only valid for amu/mmu tolerances
isotope_error = 2                      # 0=off, 1=0/1 (C13 error), 2=0/1/2, 3=0/1/2/3, 4=-1/0/1/2/3, 5=-1/0/1

#
# search enzyme
#
search_enzyme_number = 1               # choose from list at end of this params file
search_enzyme2_number = 0              # second enzyme; set to 0 if no second enzyme
sample_enzyme_number = 1               # specifies the sample enzyme which is possibly different than the one applied to the
↪ search;
                                       # used by PeptideProphet to calculate NTT & NMC in pepXML output (default=1 for
↪ trypsin).
num_enzyme_termini = 2                 # 1 (semi-digested), 2 (fully digested, default), 8 C-term unspecific , 9 N-term
↪ unspecific
allowed_missed_cleavage = 2            # maximum value is 5; for enzyme search

#
# Up to 15 variable_mod entries are supported for a standard search; manually add additional entries as needed
# format:  <mass> <residues> <0=variable/else binary> <max_mods_per_peptide> <term_distance> <n/c-term> <required>
↪ <neutral_loss>
#     e.g. 79.966331 STY 0 3 -1 0 0 97.976896
#
variable_mod01 = 15.9949 M 0 3 -1 0 0 0.0
variable_mod02 = 0.0 X 0 3 -1 0 0 0.0
variable_mod03 = 0.0 X 0 3 -1 0 0 0.0
variable_mod04 = 0.0 X 0 3 -1 0 0 0.0
variable_mod05 = 0.0 X 0 3 -1 0 0 0.0
max_variable_mods_in_peptide = 3
require_variable_mod = 0

#
# fragment ions
#
# ion trap ms/ms:  1.0005 tolerance, 0.4 offset (mono masses), theoretical_fragment_ions = 1
# high res ms/ms:    0.02 tolerance, 0.0 offset (mono masses), theoretical_fragment_ions = 0, spectrum_batch_size = 15000
#
fragment_bin_tol = 1.0005                # binning to use on fragment ions
fragment_bin_offset = 0.4              # offset position to start the binning (0.0 to 1.0)
theoretical_fragment_ions = 1          # 0=use flanking peaks, 1=M peak only
use_A_ions = 0
use_B_ions = 1
use_C_ions = 0
use_X_ions = 0
use_Y_ions = 1
use_Z_ions = 0
use_Z1_ions = 0
use_NL_ions = 0                        # 0=no, 1=yes to consider NH3/H2O neutral loss peaks

#
# output
#
```

```
output_sqtfile = 0                      # 0=no, 1=yes  write sqt file
output_txtfile = 1                      # 0=no, 1=yes, 2=Crux-formatted  write tab-delimited txt file
output_pepxmlfile = 0                   # 0=no, 1=yes  write pepXML file
output_mzidentmlfile = 0                # 0=no, 1=yes  write mzIdentML file
output_percolatorfile = 0               # 0=no, 1=yes  write Percolator pin file
num_output_lines = 1                    # num peptide results to show

#
# mzXML/mzML/raw file parameters
#
scan_range = 0 0                        # start and end scan range to search; either entry can be set independently
precursor_charge = 0 0                  # precursor charge range to analyze; does not override any existing charge; 0 as 1st
↪ entry ignores parameter
override_charge = 0                     # 0=no, 1=override precursor charge states, 2=ignore precursor charges outside
↪ precursor_charge range, 3=see online
ms_level = 2                            # MS level to analyze, valid are levels 2 (default) or 3
activation_method = ALL                 # activation method; used if activation method set; allowed ALL, CID, ECD, ETD,
↪ ETD+SA, PQD, HCD, IRMPD, SID

#
# misc parameters
#
digest_mass_range = 700.0 5000.0       # MH+ peptide mass range to analyze
peptide_length_range = 7 50             # minimum and maximum peptide length to analyze (default min 1 to allowed max 51)
max_duplicate_proteins = 10             # maximum number of additional duplicate protein names to report for each peptide ID;
↪ -1 reports all duplicates
max_fragment_charge = 3                 # set maximum fragment charge state to analyze (allowed max 5)
min_precursor_charge = 1                # set minimum precursor charge state to analyze (1 if missing)
max_precursor_charge = 6                # set maximum precursor charge state to analyze (allowed max 9)
clip_nterm_methionine = 0               # 0=leave protein sequences as-is; 1=also consider sequence w/o N-term methionine
spectrum_batch_size = 75000             # max. # of spectra to search at a time; 0 to search the entire scan range in one loop
decoy_prefix = DECOY_                   # decoy entries are denoted by this string which is pre-pended to each protein
↪ accession
equal_I_and_L = 1                       # 0=treat I and L as different; 1=treat I and L as same
mass_offsets =                          # one or more mass offsets to search (values substracted from deconvoluted precursor
↪ mass)

#
# spectral processing
#
minimum_peaks = 10                      # required minimum number of peaks in spectrum to search (default 10)
minimum_intensity = 0                  # minimum intensity value to read in
remove_precursor_peak = 0               # 0=no, 1=yes, 2=all charge reduced precursor peaks (for ETD), 3=phosphate neutral
↪ loss peaks
remove_precursor_tolerance = 1.5       # +- Da tolerance for precursor removal
clear_mz_range = 0.0 0.0                # clear out all peaks in the specified m/z range e.g. remove reporter ion region of
↪ TMT spectra
percentage_base_peak = 0.0              # specify a percentage (e.g. "0.05" for 5%) of the base peak intensity as a minimum
↪ intensity threshold

#
# static modifications
#
add_Cterm_peptide = 0.0
add_Nterm_peptide = 304.207146
add_Cterm_protein = 0.0
add_Nterm_protein = 0.0

add_G_glycine = 0.0000                  # added to G - avg.  57.0513, mono.  57.02146
add_A_alanine = 0.0000                  # added to A - avg.  71.0779, mono.  71.03711
add_S_serine = 0.0000                   # added to S - avg.  87.0773, mono.  87.03203
add_P_proline = 0.0000                  # added to P - avg.  97.1152, mono.  97.05276
add_V_valine = 0.0000                   # added to V - avg.  99.1311, mono.  99.06841
add_T_threonine = 0.0000                # added to T - avg. 101.1038, mono. 101.04768
add_C_cysteine = 57.021464              # added to C - avg. 103.1429, mono. 103.00918
add_L_leucine = 0.0000                  # added to L - avg. 113.1576, mono. 113.08406
add_I_isoleucine = 0.0000               # added to I - avg. 113.1576, mono. 113.08406
add_N_asparagine = 0.0000               # added to N - avg. 114.1026, mono. 114.04293
add_D_aspartic_acid = 0.0000            # added to D - avg. 115.0874, mono. 115.02694
add_Q_glutamine = 0.0000                # added to Q - avg. 128.1292, mono. 128.05858
add_K_lysine = 304.207146                   # added to K - avg. 128.1723, mono. 128.09496
add_E_glutamic_acid = 0.0000            # added to E - avg. 129.1140, mono. 129.04259
add_M_methionine = 0.0000               # added to M - avg. 131.1961, mono. 131.04048
add_H_histidine = 0.0000                # added to H - avg. 137.1393, mono. 137.05891
add_F_phenylalanine = 0.0000            # added to F - avg. 147.1739, mono. 147.06841
add_U_selenocysteine = 0.0000           # added to U - avg. 150.0379, mono. 150.95363
add_R_arginine = 0.0000                 # added to R - avg. 156.1857, mono. 156.10111
add_Y_tyrosine = 0.0000                 # added to Y - avg. 163.0633, mono. 163.06333
add_W_tryptophan = 0.0000               # added to W - avg. 186.0793, mono. 186.07931
add_O_pyrrolysine = 0.0000              # added to O - avg. 237.2982, mono  237.14773
add_B_user_amino_acid = 0.0000          # added to B - avg.   0.0000, mono.   0.00000
```

```
add_J_user_amino_acid = 0.0000         # added to J - avg.   0.0000, mono.   0.00000
add_X_user_amino_acid = 0.0000         # added to X - avg.   0.0000, mono.   0.00000
add_Z_user_amino_acid = 0.0000         # added to Z - avg.   0.0000, mono.   0.00000

fragment_index_predicted_mask_file =   # Carafe predicted-fragment mask (.fi_mask); empty = disabled. Set by run_rts.ps1 via
↪ RealtimeSearch.exe --mask; kept empty here for the .idx build
carafe_mask_min_relative_intensity = 0.10   # read by tools/carafe.py prerun --params only
carafe_mask_min_peaks = 6                   # read by tools/carafe.py prerun --params only
#
# COMET_ENZYME_INFO _must_ be at the end of this parameters file
# Enzyme entries can be added/deleted/edited

#
[COMET_ENZYME_INFO]
0.  Cut_everywhere         0      -           -
1.  Trypsin                1      KR          P
2.  Trypsin/P              1      KR          -
3.  Lys_C                  1      K           P
4.  Lys_N                  0      K           -
5.  Arg_C                  1      R           P
6.  Asp_N                  0      DN          -
7.  CNBr                   1      M           -
8.  Asp-N_ambic            1      DE          -
9.  PepsinA                1      FL          -
10. Chymotrypsin           1      FWYL        P
11. No_cut                 1      @           @
```
